# The homodimeric structure of the LARGE1 dual glycosyltransferase

**DOI:** 10.1101/2022.05.11.491581

**Authors:** Michael Katz, Ron Diskin

**Affiliations:** Department of Chemical and Structural Biology, Weizmann Institute of Science, Rehovot 7610001, Israel

## Abstract

LARGE1 is a bifunctional glycosyltransferase responsible for generating a long linear polysaccharide termed matriglycan that links the cytoskeleton and the extracellular matrix and is required for proper muscle function. This matriglycan polymer is made with an alternating pattern of xylose and glucuronic acid monomers. Mutations in the LARGE1 gene have been shown to cause life-threatening dystroglycanopathies through the inhibition of matriglycan synthesis. Despite its major role in muscle maintenance, the structure of the LARGE1 enzyme and how it assembles in the Golgi are unknown. Here we present the structure of LARGE1, obtained by a combination of X-ray crystallography and single-particle cryo-EM. We found that LARGE1 homo-dimerizes in a configuration that is dictated by its coiled-coil stem domain. The structure shows that this enzyme has two canonical GT-A folds with each of its catalytic domains. In the context of its dimeric structure, the two types of catalytic domains are brought into close proximity from opposing monomers to allow efficient shuttling of the substrate between the two domains. Together with putative retention of matriglycan by electrostatic interactions, this dimeric organization offers a possible mechanism for the high processivity of LARGE1. The structural information further reveals the mechanisms in which disease-causing mutations disrupt the activity of LARGE1. Collectively, these data shed light on how matriglycan is synthesized alongside the functional significance of glycosyltransferase oligomerization.

## Introduction

Like-acetylglucosaminyltransferase 1 (LARGE1) is a bifunctional glycosyltransferase (GTase) [1] that plays a critical role in maintaining proper muscle function [2]. This enzyme is responsible for a post-translational modification of the extracellular matrix receptor, α-dystroglycan (α-DG), which enables it to interact with proteins containing a laminin-G domain [3, 4]. This carbohydrate addition to α-DG is critical for the communication between the F-actin cytoskeleton network and the basal lamina [5, 6]. The polysaccharide which LARGE1 synthesizes, termed matriglycan, is composed of xylose (Xyl) and glucuronic acid (GlcA) subunits in a linear repeating structure of [-3GlcA-β1,3-Xyl-α1-]_n_ [7, 8]. Matriglycan is thought to be a unique modification of α-DG [7] and the initiation of its synthesis depends on a complex network of at least 16 other enzymes [9-12]. The length of α-DG-conjugated matriglycan chains varies substantially and is highly dependent on the tissue where it is produced [6, 7].

Mutations in the genes responsible for the synthesis of these glycan precursors can result in hypoglycosylation of α-DG and disease-causing phenotypes [13]. Improper glycosylation of α-DG is responsible for various forms of congenital muscular dystrophy (CMD), collectively termed dystroglycanopathies, which yield skeletal, brain, and eye abnormalities [14, 15]. Mutations specifically within the *LARGE1* gene can cause serious forms of secondary dystroglycanopathies [16, 17]. Walker-Warburg Syndrome (WWS) is considered to be the most severe form of CMD and is characterized by significant brain abnormalities, inability to develop significant motor function, and typically death within infancy [14, 15]. Several point mutations within LARGE1 have been known to result in life-threatening phenotypes [15, 17-19]. *LARGE1* deficiency is also believed to be an underlying cause of epithelium-derived [20] & lung [21] cancers.

LARGE1 is present in the Golgi [22] as a type II membrane protein, composed of a transmembrane/cytosolic region, a coiled-coil/stem (CC), and domains with xylosyltransferase (Xyl-T) and glucuronyltransferase (GlcA-T) catalytic activities, which are separated by a linker [1] (Fig. 1a). Most of the currently available structural data for GTases were derived using X-ray crystallography. However, the organization of the GTases solved within a crystal lattice could make it difficult to determine their true native oligomeric states. Structural data on bifunctional GTases, like LARGE1, is scarce [23-26]. While the presence of the CC domain in LARGE1 suggests that it forms higher-order oligomers, it is unknown how LARGE1 would oligomerize. Additionally, how the two domains orchestrate alternating synthesis of this polysaccharide is unclear. The vast majority of GTases do oligomerize and the oligomeric state can be required for the enzymatic function [27, 28]. It was previously suggested that the transmembrane/stem region of a GTase is critical for their oligomerization [29, 30] but it is still unclear as to which other factors facilitate oligomer formation.

**Fig. 1.**
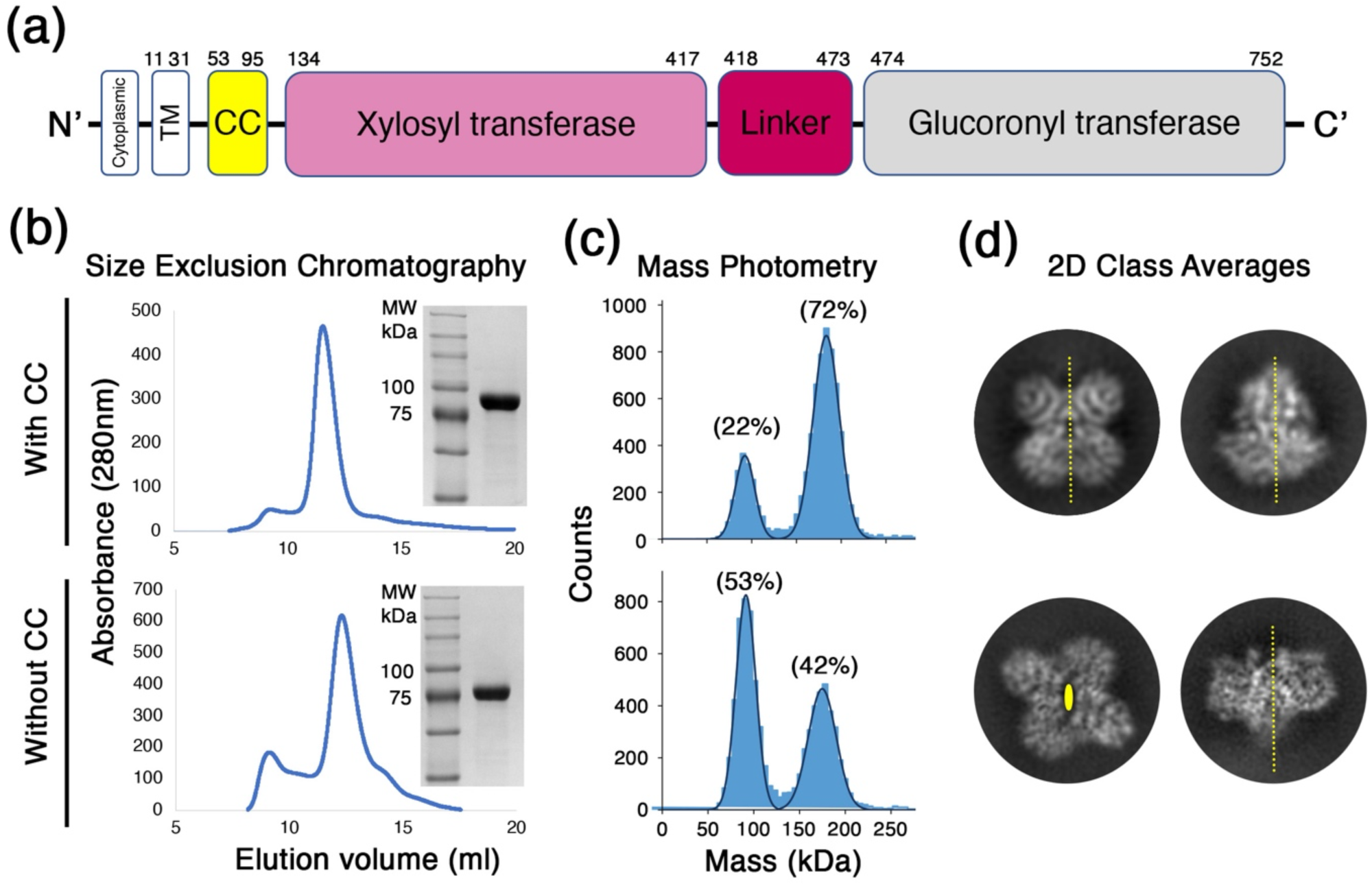
LARGE1 dimerizes in solution. (a) A schematic diagram for the domain organization in the LARGE1 protein. (b) Purification of the recombinant LARGE1 proteins. Chromatograms from SEC for constructs with (upper image) or without (lower image) the CC domain. Insets show Coomassie-stained SDS-PAGE for the purified proteins. (c) Mass-photometry analysis of the two LARGE1 proteins at a concentration of 30 nM. The fraction of counts is noted for the two main peaks in the data sets that corresponds to the monomeric (∼80 kDa) and dimeric (∼160 kDa) forms of LARGE1. The rest of the counts (6% and 5%, for with and without CC, respectively) do not belong to these two main peaks. (d) 2D class averages of LARGE1 particles from the two forms. For each LARGE1 form, two separate classes are shown, representing two distinct and perpendicular views. The 2-fold symmetry axes are shown as dashed yellow lines (for in-plane axes) or as an oval shape (for out-of-plane axis).

Here, we investigated the structure of LARGE1 using a combination of X-ray crystallography and single-particle cryo-electron microscopy (cryo-EM). We found that in solution, the catalytic domains of LARGE1 can dimerize in two distinct configurations. We further show that the CC domain promotes dimerization and further selects one of these configurations, which places the active sites of one Xyl-T and one GlcA-T in close proximity that may promote efficient synthesis of matriglycan.

## Results

### The ectodomain of LARGE1 forms a dimer in solution

To characterize LARGE1, we produced in HEK293F cells two soluble, affinity-tagged, secreted forms of this protein. Both forms include the entire catalytic ectodomain module without the transmembrane part of the protein, but one contains the CC domain, and the other does not. Both forms were readily expressed and were easily captured from cells media using Ni^2^+ affinity chromatography. Subsequent purification of these two proteins using size exclusion chromatography (SEC) resulted in nearly-homogenous protein samples (Fig. 1b). For both protein samples, the SEC elution profiles were similar, with a single main peak that contained the LARGE1 protein (Fig. 1b). Analyzing both protein samples using mass photometry at a concentration of 30 nM (2.5 μg/ml) indicated the existence of a monomer/dimer equilibrium (Fig. 1c). At this concentration, the LARGE1 with the CC domain had a higher propensity to form a dimer compared with the version of LARGE1 that did not have the CC domain (Fig. 1c). Nevertheless, even in the absence of the CC domain, the catalytic module of LARGE1 is sufficient to mediate dimerization.

While both LARGE1 constructs dimerize in solution, single-particle cryo-EM analysis indicated that they do so in two different configurations: 2D class averages indicate that in the presence of the CC domain, the dimer forms with a parallel orientation in which each of the two catalytic domains (Fig. 1a) interacts with the same domain of the second protomer (Fig. 1d). In contrast, the LARGE1 construct that is missing the CC domain forms an anti-parallel dimer in which each of the catalytic domains interacts with the other domain of the second protomer (Fig. 1d). Therefore, the ectodomain of LARGE1 has an inherent propensity to dimerize and the presence of the CC domain favors a parallel orientation for dimer formation.

### The dimeric structure of LARGE1

Single-particle cryo-EM analysis of the two LARGE1 constructs yielded near atomic-resolution density maps (Fig. 2). Both dimers exhibit a clear 2-fold symmetry, which was applied during reconstruction. The LARGE1 dimers with and without the CC domain provided density maps extending to gold-standard Fourier shell correlation (FSC) resolution values of 3.9 Å, and 3.7 Å, respectively. While obtaining the single-particle EM data, we also managed to crystalize the LARGE1 construct that lacked the CC domain and to collect X-ray diffraction data, extending to 2.6 Å (Table 1). We, therefore, utilized one-half of the EM density map (Fig. 2) as a search model in Phaser [31] to obtain phases for the X-ray diffraction data, following a previously described method [32]. The crystal of LARGE1 belonged to a *C*121 space group, and we found two copies of LARGE1 in the asymmetric unit. We used Phenix-Autobuild [33] to provide an initial model and then manually completed the model using an iterative model building and refinement in Coot [34] and Phenix-Refine [35], respectively.

**Table 1.** Data collection and refinement statistics. Statistics for the highest-resolution shell are shown in parentheses.

|  |  |
| --- | --- |
|  | LARGE1 - CC* |
| Wavelength | 0.9677 |
| Resolution range | 45.64 - 2.61 (2.76 - 2.61) |
| Space group | *C 1 2 1* |
| Unit cell | 196.531 107.377 100.334 90 120.26 90 |
| Total reflections | 140137 |
| Unique reflections | 91901 |
| Multiplicity | 1.52 |
| Completeness (%) | 84.6 (83.3) |
| Mean I/sigma(I) | 3.57 (0.35) |
| Wilson B-factor | 77.82 |
| R-meas | 17.7 (263.6) |
| CC<sub>1/2</sub> | 98.8 (11.7) |
| Reflections used in refinement | 91817 |
| Reflections used for R-free | 2116 |
| R-work | 0.231 |
| R-free | 0.288 |
| Number of non-hydrogen atoms | 9734 |
| macromolecules | 9685 |
| ligands | 45 |
| solvent | 4 |
| Protein residues | 1170 |
| RMS(bonds) | 0.005 |
| RMS(angles) | 0.91 |
| Ramachandran favored (%) | 97.38 |
| Ramachandran allowed (%) | 2.62 |
| Ramachandran outliers (%) | 0.00 |
| Rotamer outliers (%) | 3.53 |
| Clashscore | 10.72 |
| Average B-factor | 100.91 |
| macromolecules | 100.8 |
| ligands | 126.38 |
| solvent | 65.71 |
| Number of TLS groups | 9 |

**Fig. 2.**
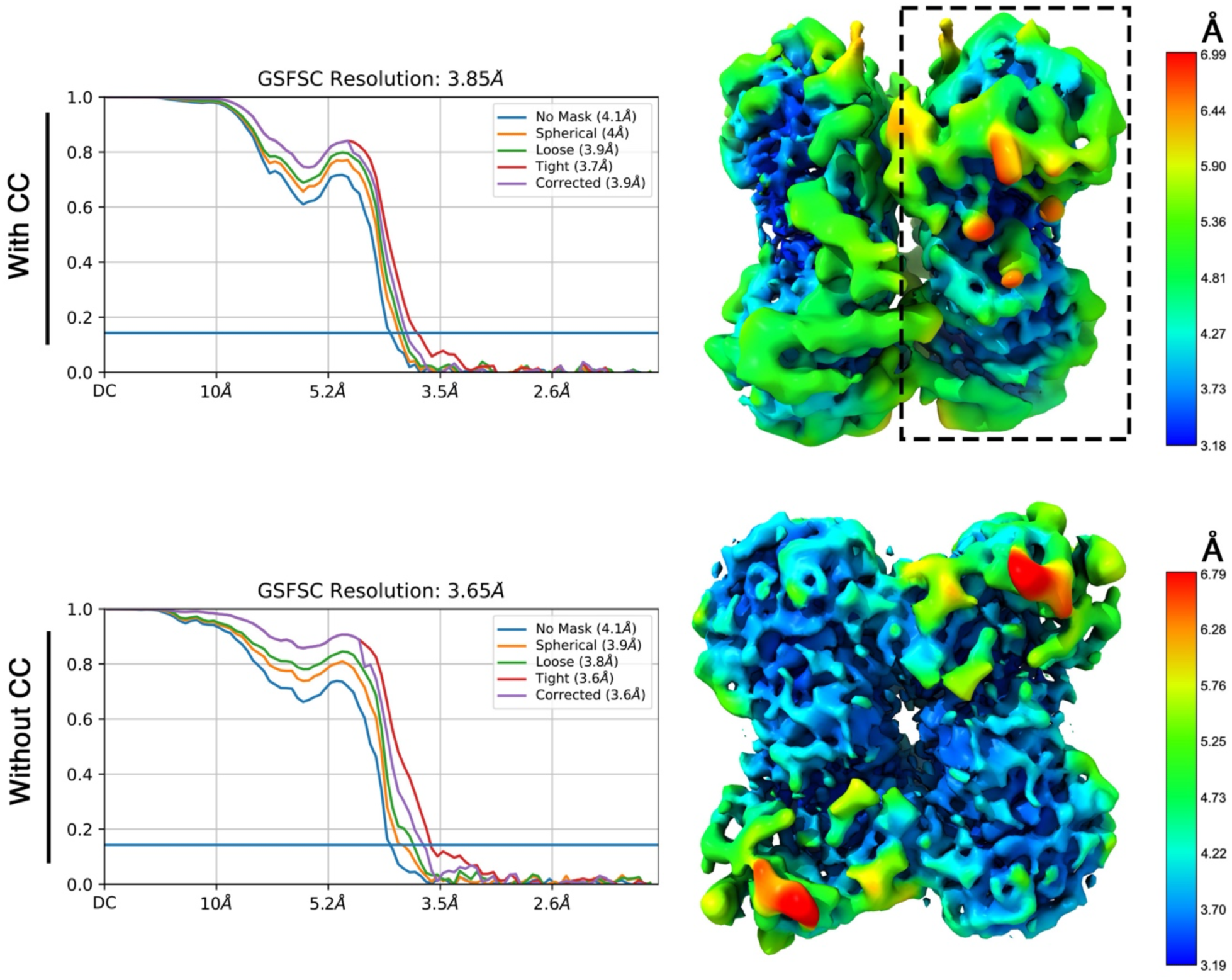
Single particle cryo-EM maps of LARGE1. Reconstructed maps of the LARGE1 constructs with the CC domain (upper part) and without the CC (lower part). The gold standard FSC curves are shown on the left and density maps colored by the local resolution estimates are shown on the right. One half map that was used as a molecular replacement model is highlighted with a dashed rectangle.

The crystal structure of LARGE1 consists of two chains that make a dimer in the asymmetric unit (Fig. 3a). For both chains (i.e., A and B), electron density was visible and hence allowed us to model residues 134 to 752. The electron density for some loops was either poor or was lacking completely and hence the model lacks residues: 282-286, 362-367, 369-372, 382-391, 626-636, 718-719 for chain A, and 280-286, 361-367, 370-372, 383-391, 713-718 for chain B. The two chains adopt an almost identical configuration with a root mean square deviation (RMSD) value of 1.4 Å for all shared Cα atoms (Fig. 3a). Despite the fact that we have crystalized the LARGE1 construct that did not have the CC domain, the two chains in the asymmetric unit formed a parallel dimer that resembles the EM map of the LARGE1 with the CC domain. Indeed, the crystallographic dimer readily fits into the EM density map of the LARGE1 with the CC domain (Fig. 3b). In order to fit the EM map of the LARGE1 without the CC domain, the chains need to be rotated in respect to one another to form an anti-parallel dimer (Fig. 3c). Considering the high similarity of chains, A and B of the crystallographic model (Fig. 3a) and since a rotation of the LARGE1 monomers is sufficient to get a good fit into the EM map of the LARGE1 without the CC domain, it indicates that the relative orientation of the Xyl-T domain in respect to the GlcA-T domain is maintained in all of these states. In addition, the fact that the LARGE1 protein without the CC domain crystalized in the parallel dimeric form as observed for the LARGE1 protein with the CC domain (Fig. 3b) implies that it also forms such parallel dimers in solution despite the fact that the EM analysis only revealed anti-parallel dimers (Figs. 1d, 2, and 3c). Hence, these low-abundant parallel dimers were the ones that nucleated and dictated the organization of the LARGE1 crystals.

**Fig. 3.**
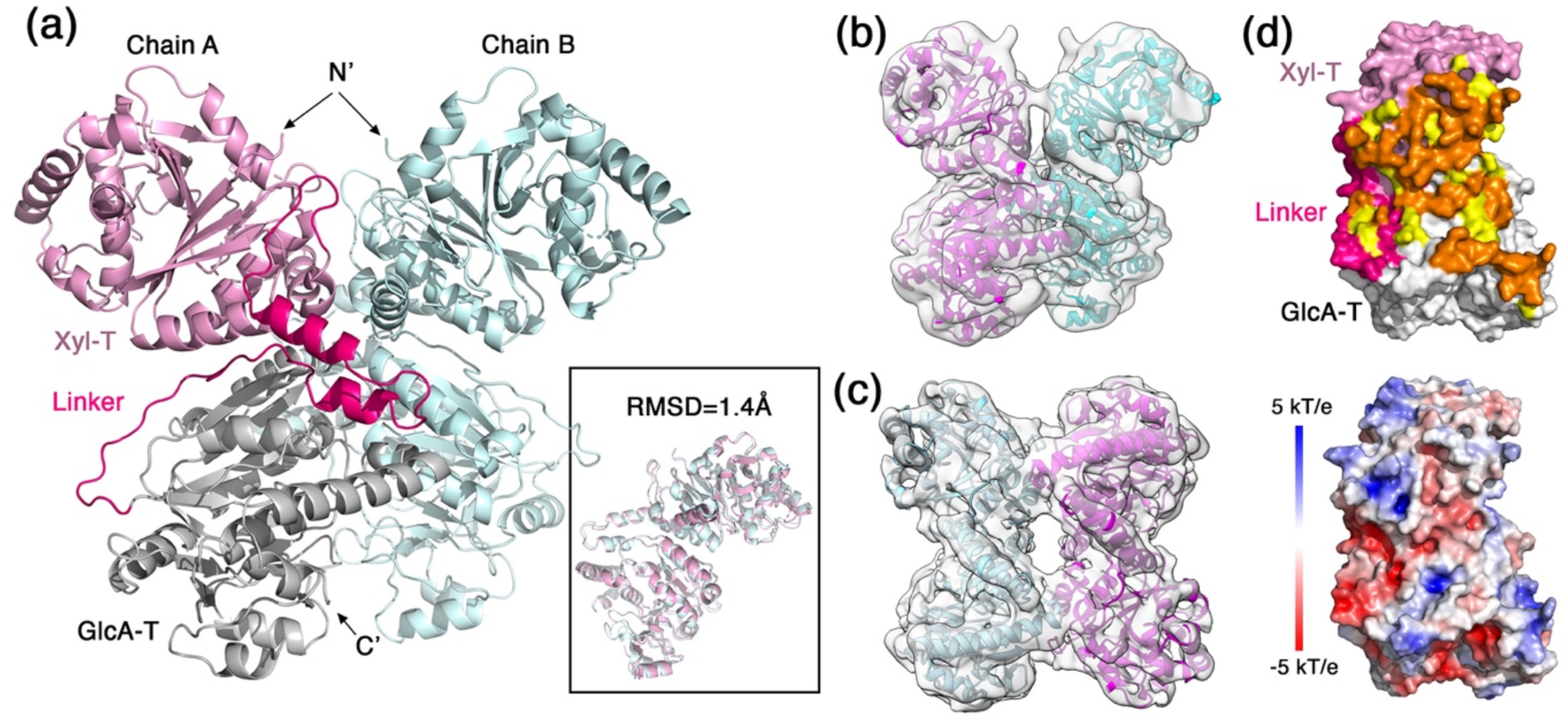
The dimeric structure of LARGE1. (a) Ribbon representation of the two LARGE1 monomers that make the asymmetric unit of the crystal. The Xyl-T, Linker region, and GlcA-T regions are highlighted in pink, hot pink, and gray on one monomer, respectively. The N- and C-termini are noted. Inset shows a superimposition of the two modeled LARGE1 chains. (b) A fit of the LARGE1 dimer as appeared in the asymmetric unit of the crystal into the EM density map of the LARGE1 with the CC domain. (c) An individual fit of two LARGE1 monomers into the EM density map of the LARGE1 protein without the CC domain. (d) Analysis of the dimer interface. Upper image shows a LARGE1 monomer in a surface representation. The Xyl-T, Linker region, and GlcA-T regions are highlighted in pink, hot pink, and gray, respectively. Fully buried surface residues due to dimerization are shown in orange (probe size of 1.4 Å). Partially buried surface residues (i.e., making cavities) due to dimerization are shown in yellow (probe size of 2.0 Å). Lower image shows a LARGE1 monomer in the same orientation as the upper image with surface representation that is color-coded by surface electrostatic potential (blue, 5 kT/e and red, -5 kT/e).

The selection of the parallel dimeric form by the presence of the CC domain (Figs. 1d, 3b) indicates that this may be the preferred physiological dimer *in-vivo*. We hence focus our analysis on this dimeric form. Inspection of the parallel dimer interface shows an overall buried surface area of 2,692 Å^2^ (1,401 Å^2^ on chain ‘A’ and 1291 Å^2^ on chain ‘B’) (Fig. 3d). However, if we consider additional residues at the interface, which make very narrow cavities (i.e., using a probe with a radius of 2.0 Å for detecting buried surfaces), the buried surface area then becomes 4,496 Å^2^. This buried surface area spans the Xyl-T and GlcA-T catalytic domains as well as some residues at the linker region between the domains (Fig. 3d). Examining the surface electrostatic potential shows that the dimer interface is mostly uncharged (Fig. 3d). A large hydrophobic surface may help to rationalize the promiscuity of LARGE1 in the geometry of dimer formation when the dimer architecture is not predefined by the CC domain.

### The Xyl-T and GlcA-T catalytic domains

The Xyl-T domain of LARGE1 has a canonical GT-A fold, and its catalytic site contains a conserved DXD motif [36]. This domain is composed of 7-stranded major & 3-stranded minor β-sheets, encompassed by α-helices. Comparing Xyl-T with the structure of a mouse-derived xyloside α-1,3-xylosyltransferase (XXYLT1) [37] reveals a similar general architecture (Fig. 4a). In the active site of the Xyl-T domain, we could not detect a clear density that would indicate the presence of a divalent cation. Nevertheless, the superimposition of the structures places the XXYLT1-derived Mn^2+^ such that it would be coordinated by Asp242, and Asp244, which make the DXD motif, and by His380 of LARGE1 (Fig. 4a). These three residues are conserved in XXYLT1 as well as in other xylosyltransferases [37, 38]. The Mn^2+^ helps to position the UDP-xylose in the active site [37]. Based on the XXYLT1 structure, we can postulate the position of the UDP-sugar donor in the active site of the LARGE1 Xyl-T (Figs. 4a, 4b). Gln335 of LARGE1 is also a conserved residue (Fig. 4a), and based on the enzymatic mechanism of XXYLT1 [37], it binds the substrate and likely also functions in stabilizing high-energy intermediate states, making it a critical residue for catalysis. Overall, the superimposition with XXYLT1 elucidates how the UDP-xylose and growing substrates would fit into the catalytic site of the LARGE1 Xyl-T (Fig. 4b).

**Fig. 4.**
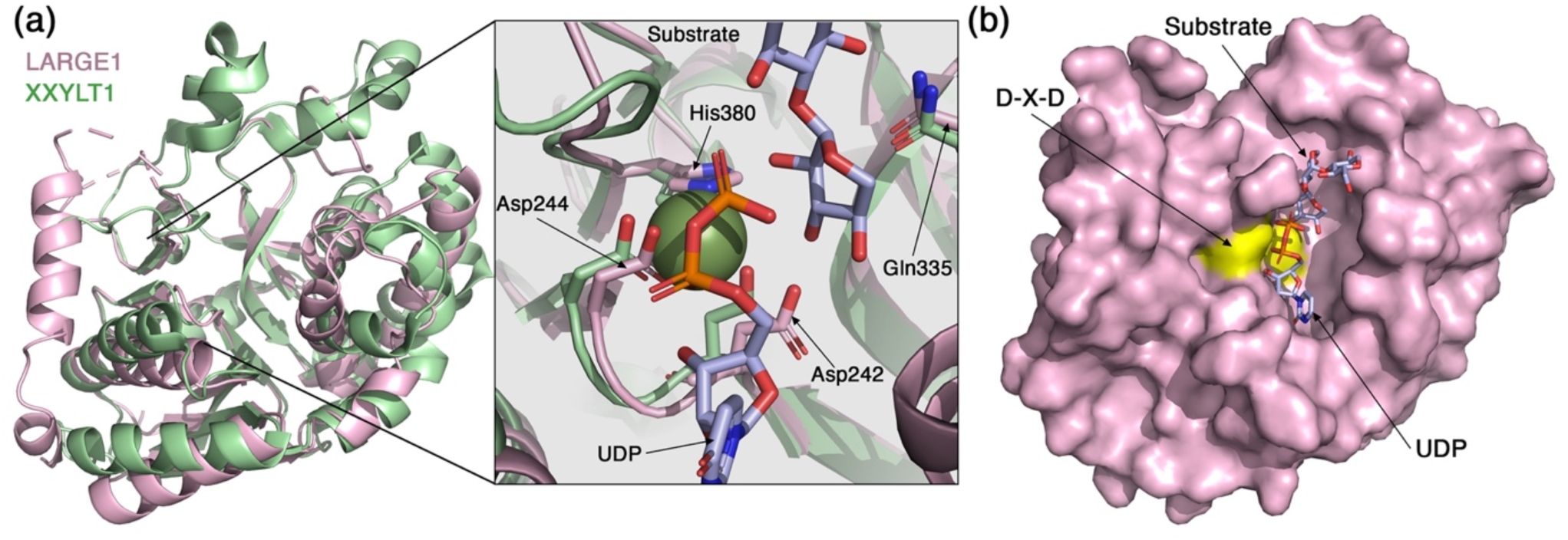
The Xyl-T domain of LARGE1. (a) Superimposition of the Xyl-T domain from LARGE1 (pink) with XXYLT1 (PDB: 4wn2, green). Inset shows a closeup view of the active site. The conserved residues that coordinate the Mn^2+^ are notes, as well as a UDP and a growing substrate that were resolved in the structure of XXYLT1. (b) Surface representation of the LARGE1 Xyl-T domain. The aspartic acids of the DXD motif are colored yellow. The UDP and the growing substrate derived from the structure of XXYLT1 are shown as sticks.

The GlcA-T domain of LARGE1 also adopts a canonical GT-A fold and has a conserved DXD motif (Fig. 5a). The closest structural homolog of the LARGE1 GlcA-T according to 3D-BLAST [39] is the chondroitin polymerase from K4 *E*.*coli* (K4CP) (PDB:2z86) [40]. Similar to LARGE1, K4CP is a bifunctional glycosyltransferase that catalyzes the addition of GlcA and *N*-acetylgalactosamine to the chondroitin polymer [40]. In the electron density map, we observed density for a coordinated ion in the active site of the GlcA-T domain (Fig. 5b) and we therefore modeled a Mn^2+^ atom. Compared with K4CP, the Mn^2+^ atoms in the GlcA-T domain are slightly shifted away from the coordinating aspartic acids and histidine residues (Fig. 5a, inset). The exact position of Mn^2+^ may change when UDP-GlcA is bound in the active site as this shifted location seems to collide with the UDP as seen in the K4CP structure. The LARGE1 GlcA-T domain has an extra histidine (His708) that seems to participate in the binding of the Mn^2+^. This histidine residue is not conserved in K4CP which has a proline (Pro389) instead (Fig. 5a, inset). As in the Xyl-T domain, the Mn^2+^ in the GlcA-T domain is important for holding the UDP-sugar donor in place. A surface representation of the GlcA-T domain reveals the approximate location of the UDP in the active site (Fig. 5c). The K4CP-derived UDP fits into a deep pocket, leaving a narrow tunnel for the growing matriglycan substrate to enter (Fig. 5c).

**Fig. 5.**
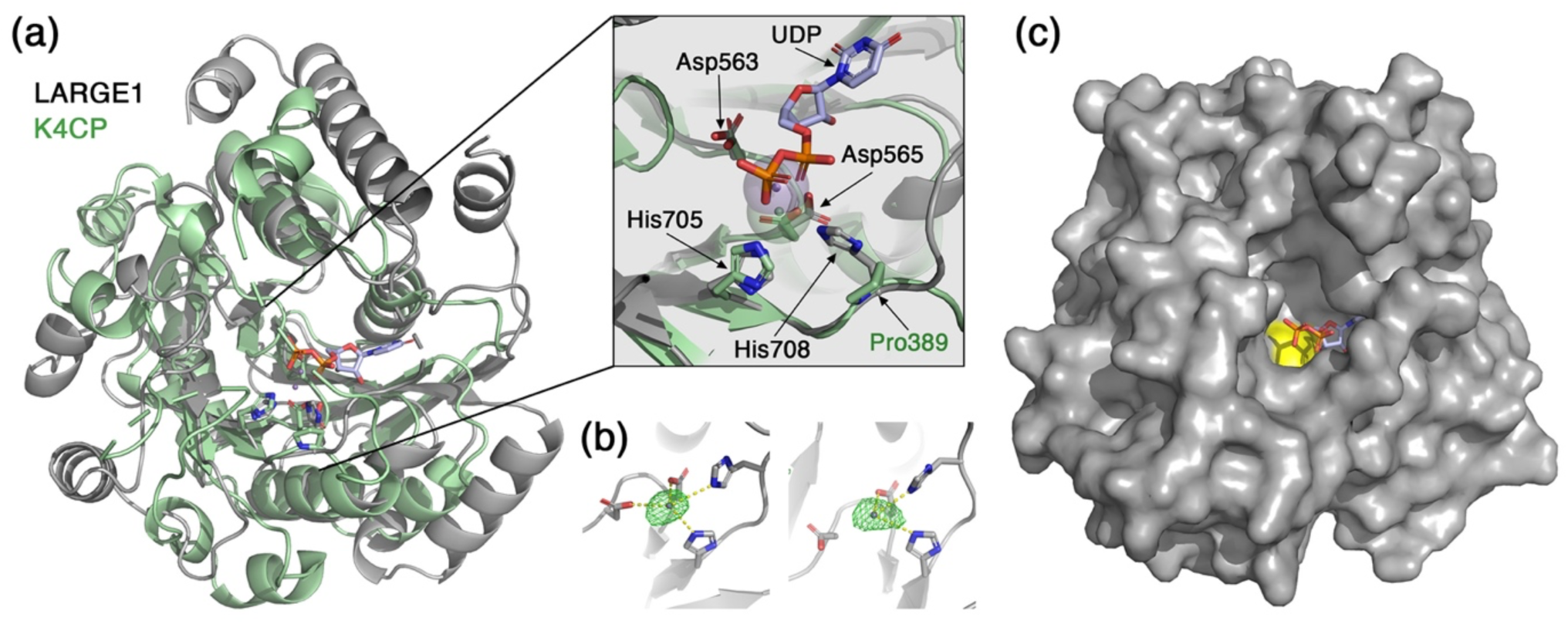
The GlcA-T domain of LARGE1. (a) Superimposition of the GlcA-T domain from LARGE1 (grey) with K4CP (PDB: 2z86, green). Inset shows a closeup view of the active site. Mn^2+^ atoms of LARGE1 (purple) and of K4CP (green) are shown as semi-transparent spheres. A UDP from the K4CP structure is shown in purple. The conserved DXD motif and histidine residues that participate in the coordination of the Mn^2+^ are noted. (b) The GlcA-T active sites of the two LARGE1 monomers (left and right) in the crystallographic asymmetric unit are shown with an Fo-Fc difference map (green mesh, σ=5) calculated after omitting the Mn^2+^ atoms. (c) Surface representation of the LARGE1 GlcA-T domain. The aspartic acids of the DXD motif are colored yellow. The UDP derived from the structure of K4CP is shown as sticks.

### Matriglycan synthesis by the dimeric LARGE1

The matriglycan polymer that LARGE1 synthesizes is made of alternating Xyl and GlcA monomers. Hence, the growing matriglycan chain needs to enter into the active sites of the Xyl-T and the GlcA-T domains in an alternating manner as well. Interestingly, the closest Xyl-T and GlcA-T sites in the context of the LARGE1 dimer are located on opposite monomers rather than on a single protomer (Fig. 6a). These two sites are approximately 40 Å away from each other and make the shortest possible path for a matriglycan non-reducing end to encounter the two different transferases. The alternative path requires the matriglycan to travel around the dimer to reach its far side. Hence, most efficient synthesis is theoretically achieved when the Xyl-T and GlcA-T active sites are contributed from two distinct monomers. Also, the matriglycan product can potentially consist of hundreds of repeating Xyl-GlcA units [7]. This implies that LARGE1 has high processivity for synthesizing matriglycan. Such high processivity could be aided if the growing matriglycan chain will not be completely released after the addition of each sugar monomer. Due to the high content of GlcA, matriglycan has a negative charge. Examining the surface electrostatic potential of the LARGE1 dimer, we see a positively-charge patch at the interface of the two GlcA-T domains and a significantly larger positively-charged surface formed at the interface of the Xyl-T domains (Fig. 6b). We speculate that unspecific electrostatic interactions may form between these positively-charged surfaces of LARGE1 and with matriglycan, which could prevent a complete release of the polymer after each step of sugar addition. By that, the effective local concentration of matriglycan will increase, which will promote synthesis and will contribute to the processivity of LARGE1.

**Fig. 6.**
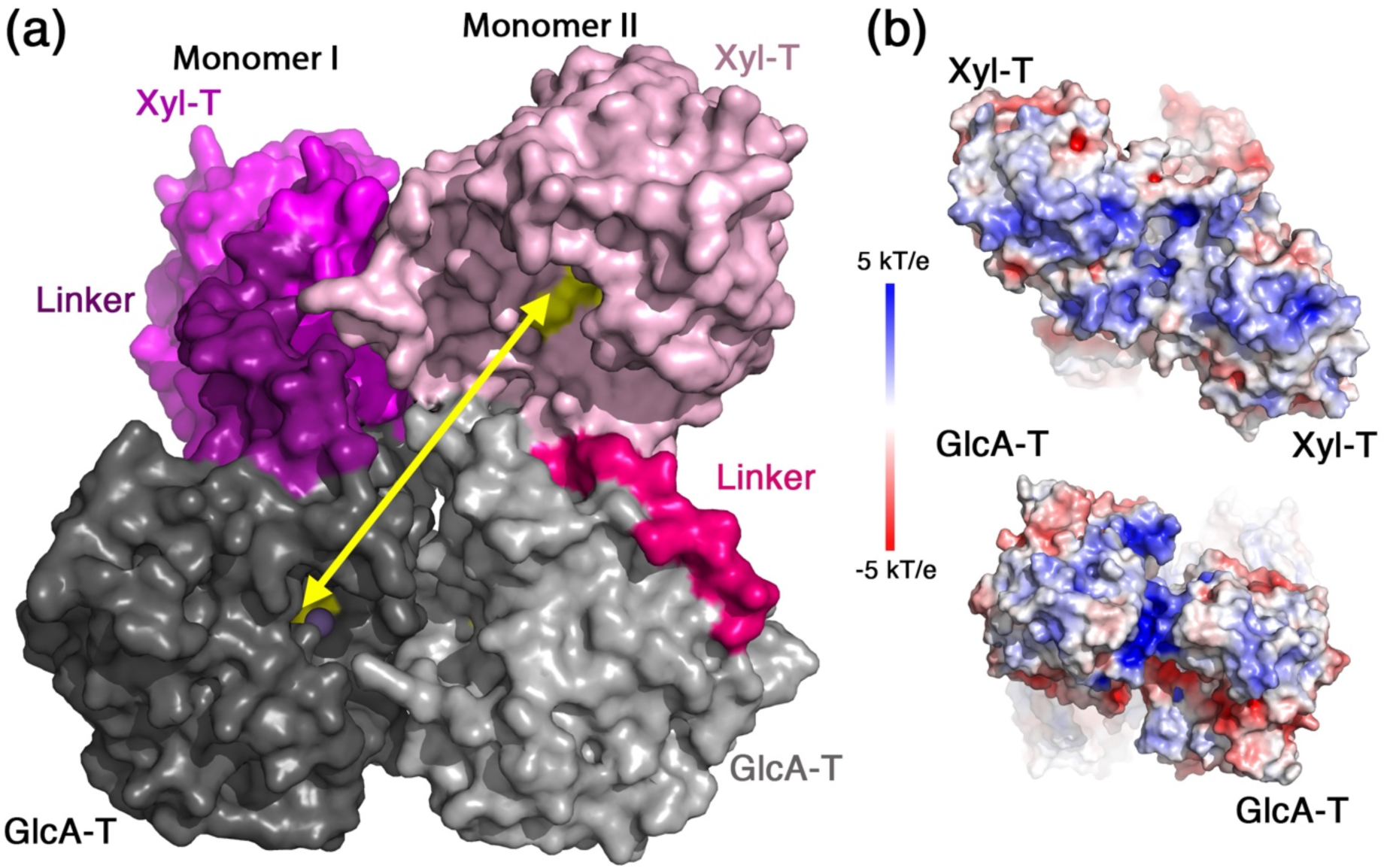
Matriglycan synthesis by the LARGE1 dimer. (a) The closest active sites of the Xyl-T and GlcA-T domains are located of different monomers. The LARGE1 dimer is shown using a surface representation and the catalytic domains are colored as in Fig. 3a. The two LARGE1 monomers are highlighted by different tones. A yellow arrow shows the openings of the closest Xyl-T and GlcA-T catalytic sites. (b) Surface electrostatic potentials of the LARGE1 dimer from the Xyl-T (top) and from the GlcA-T (bottom) sides of the dimer. Blue to red colors indicate 5 kT/e to -5 kT/e, respectively.

### Disease-causing mutations in LARGE1

Several alterations in LARGE1 were previously identified as disease-causing mutations. Such mutations include: S331F [18], C443Y [15], W495R [19], E509K [17]. Having the structural information for LARGE1 allows us to gain some insights for the possible molecular mechanisms that underly the deleterious effect of these mutations. Ser331 is a surface-exposed residue in the Xyl-T catalytic domain (Fig. 7a). Introducing phenylalanine in this position does not have an obvious consequence on the structure of LARGE1. Nevertheless, Ser331 is located at the vicinity of some surface-exposed hydrophobic residues like Trp276 and Leu332 (Fig. 7a). Potentially, introducing a S331F mutation may form a hydrophobic patch that would drive aggregation of LARGE1. Cys443 is located at the linker between the Xyl-T and GlcA-T domains (Fig. 7a). It makes a disulfide bond with a cysteine residue at the GlcA-T domain (Fig. 7a). A tyrosine residue instead of a cysteine in position 443 will abrogate the disulfide bridge and will likely also prevent proper folding due to steric clashes with its bulky side chain. Trp495 is located at the GlcA-T domain and is making an integral part of the hydrophobic core (Fig. 7a). A positively-charge arginine facing straight into the hydrophobic core of the protein is likely to be highly destabilizing for the folded state of the protein, which will effectively prevent its proper folding. Glu509 is located at the dimer interface of the GlcA-T domains near its 2-fold symmetry axis (Fig. 7a). Glu509 is making long-range electrostatic interaction with a nearby Lys536 (Fig. 7a), which likely balance the overall surface electrostatic potential at this position. Mutating Glu509 to a lysine will likely cause the accumulation of a local positive surface electrostatic potential, which will make dimer formation to be less favorable.

**Fig. 7.**
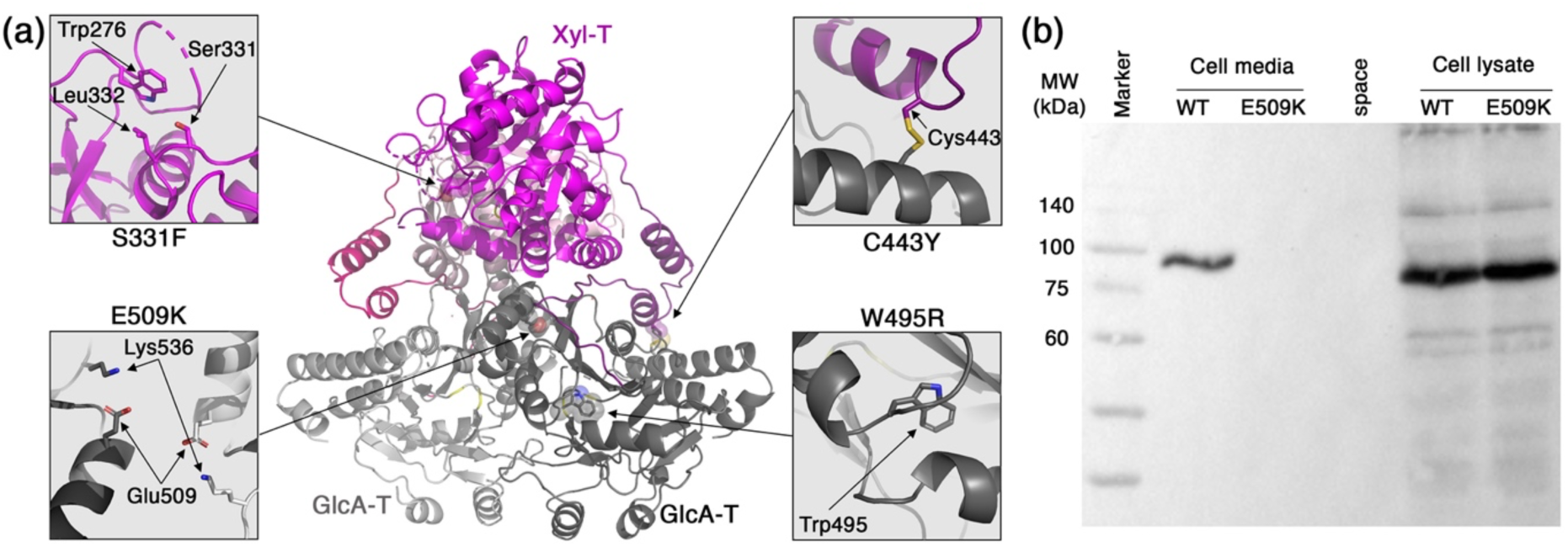
Disease causing mutations in LARGE1. (a) Mapping of four mutations on the structure of LARGE1. Insets show closeup views of the native residues at the mutation sites. (b) Western blot analysis using anti-His antibody of the production of soluble LARGE1-His WT or E509K mutant. Samples include the total cell lysates and cells’ media as indicated.

From the abovementioned structural analysis, it seems like the underlying molecular mechanisms of these four mutations involve the disruption of the global structure of LARGE1 rather than a direct interference at the catalytic sites. Indeed, after individually incorporating each of the four point mutations on the secreted form of LARGE1 that has the CC domain, we could not detect secreted LARGE1 in the cells’ media. A more complete analysis of one of the mutations, *i*.*e*., E509K, indicates that the protein is actually produced in cells but fails to secrete (Fig. 7b). This is likely due to the quality control system of the endoplasmic reticulum [41], which supports the notion of a global effect of these mutations on the stability of LARGE1.

## Discussion

LARGE1 has a high propensity to form dimers. Mass-photometry measurements at LARGE1 concentration of 30 nM indicated the presence of significant amounts of both the monomer and of the dimer forms in equilibrium (Fig. 1c). At higher concentrations, as were used for preparing the EM grids for example (*i*.*e*., ∼4 μM), the predominant form was already dimeric as no 2D class averages were obtained for a monomeric LARGE1. It is also clear that the CC domain promotes dimerization of LARGE1 (Fig. 1c) and further selects a parallel organization for the dimer (Figs. 1d, 2, & 3b). Taken together, under physiological conditions, in cells, LARGE1 is most likely in a form of a parallel dimer. Interestingly, this parallel dimer form of LARGE1 includes homo-association of the Xyl-T and the GlcA-T domains (Fig. 3a). Homo-association of other Xyl-Ts and GlcA-Ts were previously observed and in some cases were found to be important for their enzymatic activities [28, 37, 42-45].

The assembly of LARGE1 into a dimer brings into a close proximity the active sites of Xyl-T and GlcA-T from opposite monomers (Fig. 6a). This proximity between the two active sites increases the probability that the growing matriglycan chain will be sequentially modified by the two domains before the matriglycan polymer dissociates from LARGE1. Furthermore, we hypothesize that unspecific electrostatic interactions between patches with positive surface electrostatic potential on LARGE1 (Fig. 6b) and the negatively-charged matriglycan may form and thus reduce the propensity of matriglycan to completely dissociate from LARGE1 during synthesis. Acting together, these properties of LARGE1 may promote its processivity and contribute to its ability to produce long matriglycan polymers.

Several point mutations within the *LARGE1* gene were linked to pathological conditions. The E509K missense mutation results in a significant loss of matriglycan generation, along with α-DG:laminin binding, and thus causes a serious form of α-dystroglycanopathy, termed MDC1D [17, 46]. The E509K LARGE1 mutant was shown to localize in the ER and does not progress to the Golgi [46]. Our observations corroborate this notion as the soluble His-tagged LARGE1 that carries the E509K mutation is also produce in cells but is not secreted (Fig. 7b), indicating a failure to progress in the secretory pathways. Our structural analysis suggests that the E509K mutation will abrogate dimerization of LARGE1 (Fig. 7a), implying that proper dimerization of LARGE1 is required for progressing through the ER/Golgi network. Interestingly, oligomerization has been shown to be a prerequisite for Golgi translocation for other membrane-bound proteins. The GTases, EXT1 & EXT2 that function as a heterodimer for example, are retained in the ER when expressed alone. Only upon co-expression of both of these GTases the heterodimer form and can reach the Golgi [47, 48]. Other membrane proteins, such as prenylin, require homodimerization to enter the Golgi [49], and a failure to multimerize retains various other proteins in the ER [50-53]. Taken together, dimerization of LARGE1 is essential for its proper function in cells.

Overall, we have deciphered the structure of the LARGE1 bi-functional glycosyltransferase. Dictated by the CC stem domain, a parallel dimer of LARGE1 is formed and is required for proper localization in the cells. Dimerization enables advantageous properties for LARGE1 that likely contribute to its ability to synthesize long chains of matriglycan.

## Materials & Methods

### Enzyme Expression & Purification

Two variants of LARGE1 were generated, both from the human LARGE1 (xylosyl- and glucuronyltransferase) gene, obtained from the Forchheimer plasmid bank (Weizmann Institute of Science, Clone ID 100000367). Both variants were cloned without the transmembrane and cytosolic tail domains into a modified pHLSEC plasmid. The cloning, expression, and purification for this enzyme have been previously described [54]. Briefly, the LARGE1 gene with its coiled-coil (+CC) domain (residues 29-756), with a His tag + GSGG linker at its C-terminal end, was cloned into *pHLsec* using BglII/NotI restriction sites. The same LARGE1 gene without its CC domain (-CC) (residues 96-276), with a 6x His tag + GSGG linker at its N-terminal end was cloned in the same manner. Both plasmids were then separately transfected into HEK293F cells at a density of approximately 1 × 10^6^ million cells/ml. The transfections occurred with 40 kDa poylethylenamine at a ratio of 1:2.5 DNA:PEI (PEI Max, PolySciences). The productions were harvested after 1 week, spun down at 600 xg (the cells were then discarded), and the supernatants was then centrifuged at 15,800 xg to remove cellular debris. The supernatant of each production was then passed through a 0.45 μm Stericup filter (Merck Millipore) and supplemented with 100 μM phenylmethyl sulfonyl fluoride (PMSF) and 0.02% sodium azide. After which, the productions were buffer exchanged to TBS (150 mM NaCl, 20 mM Tris pH 8.0) using a Pellicon Tangential Flow Filtration system (Merck Millipore) and purified by their 6x His tag by a 5 ml HiTrap IMAC FF Ni^2+^ column (GE Healthcare). Elution from the column was then carried out with 10% imidiazole, and the proteins were concentrated using a 4 ml 30 kDa Amicon (Merck Millipore) and run on a Supderdex 200 10/300 GL column (GE Healthcare) with a running buffer of TBS + 0.2% sodium azide. The aliquots from the main HPLC peak were then pooled, concentrated using another 4 ml 30 kDa Amicon (Merck Millipore), and finally flash frozen and stored at -80 °C.

### Cryo-electron microscopy data collection & analysis

LARGE1 variants (0.3 mg/ml) were plunged onto R 0.6/1 Cu/C Quantifoil grids (EMS) using a Vitrobot system (Thermo Fisher/FEI; 4 °C, 100% humidity) and then stored under cryogenic conditions. Plasma treatment for the grids was carried out at 90 seconds, 15 mA, and plunging used 3.5 μl of samples with a blot force of -1 for 3 seconds. Grids were first screened on the Talos Arctica microscope (200 kV; Thermo Scientific) and then transferred to the Titan Krios 3Gi (300 kV; FEI) with a Gatan K3 direct detection camera for data collection. The beam size used was 900 nm and the magnification was 120,000 X, with a defocus range of -1.6 to 0.6 μm and a pixel size of 0.52 Å.

Data processing was carried out using the cryoSPARC v2 software. For both LARGE1 variants, patch motion correction & CTF estimation were first completed. For the LARGE + CC variant, a total of 7,969 movies were initially obtained and 5,451 movies were then accepted for particle selection using the blob picker. Initially, 1,760,546 particles were chosen. We then extracted the particles using a 400-pixel box, which was Fourier-cropped to a 200-pixel box size with a pixel size of 1.04 Å. After multiple iterations of the 2D classifications, 92,093 particles were finally obtained which belonged to a single class. To improve the GSFCS resolution, we applied a twofold symmetry onto the LARGE1 + CC dimer. The 3D reconstruction had a final GSFSC resolution of 3.64 Å. For LARGE1 – CC (lacking the coiled-coil domain), the same data processing method was used. A total of 5,250 movies were initially obtained, and 4,896 movies were used for further processing. After patch motion correction & CTF estimation, the blob picker in cryoSPARC v2 selected 223,954 particles. Using twofold symmetry, 164,336 particles were chosen, all of which related to a single class. An electron density map of LARGE – CC with a final GSFSC resolution of 3.86 Å was then obtained.

### Crystallization and data collection

The initial crystallization hit occurred with the SaltRx screen (Hampton Research) using the Mosquito robot (LLP Labtech) in a 96-well plate (LLP). Optimization was carried out with the Dragonfly robot (LLP), where LARGE1-CC crystals were obtained in a 200 ul drop, at a 1:1 ratio (protein: crystallization reagents), containing 1.0 M Na_3_PO_4_ monobasic monohydrate, K_3_PO_4_ dibasic / pH 5.0. PEG 200 (20% final concentration) was added to the crystals as a cryopreservative before storing in liquid N_2_ for transport. Data collection was carried out at the European Synchrotron Research Facility (ESRF), with Beamline ID ID30a. A Dectris Eiger 4M detector was used. Diffraction data was successfully obtained using a wavelength of 0.9677 Å to a resolution of 2.6 Å.

### 3D Model generation of LARGE1 variants

The monomer of the cryoEM electron density maps was used in combination with the diffraction data for molecular replacement in Phenix. A model for the crystallographic structure was then prepared with Coot, after applying the Autobuild function to generate an initial model. Numerous refinement cycles for model building were then carried out. After all ligands, glycans, and additional water molecules were identified and a model was the highest completeness was prepared, the monomer of this dimeric structure was docked into each cryo-EM map to obtain the two structures (85% completeness) with and without the coiled-coil domain. Additional refinement cycles using Phenix was then carried out to obtain the final model for each variant.

### Mass Photometry

LARGE1 samples were analyzed on a Refeyn One^MP^ (Refeyne) mass photometer. The calibration curve was carried out using jack bean urease (100 μg/ml) (Sigma). LARGE1 samples were diluted in TBS to approximately 30 nM (2.5 μg/ml) and were analyzed at room temperature over 60 second recordings using their Acquire^MP^ software, at 100 frames per second (fps). Analysis occurred using Refeyne’s Discover^MP^ software.

### Mutagenesis

Mutants were produced on the LARGE1 construct containing the coiled coil in a modified pHLsec plasmid. Point mutations were carried out via the QuikChange Site-Directed Mutagenesis protocol (Aligent). After sequence verification, mutants were amplified (via MaxiPrep) and transfected into HEK293F cells.

### Western Blot analysis

Western Blot analysis was carried out using an anti-His antibody (Qiagen) at a 1:2,000 dilution, in Tris-Buffer Saline + Tween (20 mM Tris, pH 8.0, 150 mM NaCl, 0.5% Tween) containing 1% BSA, incubated overnight at 4 °C. Then, the membrane was incubated with HRP-conjugated anti-mouse secondary antibody (Jackson ImmunoResearch), diluted at 1:10,000, before being developed with EZ-ECl (Biological Industries).

## Acknowledgements

We thank Moshe Goldsmith in the Department of Biomolecular Sciences at the Weizmann Institute of Science and Adar Sonn-Segev from Refeyn Ltd. For assisting us to perform and analyze the mass photometry measurements. The Diskin lab is supported by research grants from the Ernst I Ascher foundation, Ben B. and Joyce E. Eisenberg Foundation, Estate of Emile Mimran, Jeanne and Joseph Nissim Center for Life Sciences Research, Dov and Ziva Rabinovich Endowed Fund for Structural Biology, Donald Rivin, Stanley and Tanya Rossby Endowment Fund, Natan Sharansky, Dr. Barry Sherman Institute for Medicinal Chemistry, as well as from the Israel Science Foundation (grants No. 3147/19 and 209/20).

## Notes

### Competing Interest Statement

The authors have declared no competing interest.

